# 3D-Printable Non-invasive Head Immobilization System for Non-Human Primates

**DOI:** 10.1101/2024.01.02.573832

**Authors:** Elia Shahbazi, Drew Nguyen, Tyler Swedan, Timothy Ma, Rosa Lafer-Sousa, Alvin Dinh, Reza Azadi, Amy M. Ryan, Arash Afraz

## Abstract

A critical contribution to understanding the primate brain is our ability to directly record, manipulate, or image the brain in awake, behaving monkeys. A challenge with carrying out these studies is that typically the monkey’s head must be fixed in place. While non-invasive head immobilization systems (NHIS) have been demonstrated to be effective, they have not fully replaced the use of surgically applied headposts. Here, we introduce a novel NHIS, with an option for voluntary engagement, for macaques using a widely available resource: the 3D printer. We designed customized 3D-printable head-immobilization masks for monkeys and a docking system using standard CT scans and user-friendly open-source software packages (FLoRIN, Blender, Python). We examined the efficacy and stability of our mask system for collecting eye fixation data by measuring trial-by-trial gaze precision and accuracy in two mask-immobilized monkeys and three headposted monkeys, including a within-subjects comparison. Our results demonstrate that monkeys stabilized by the mask maintain gaze precision and accuracy comparable with their headposted counterparts. While the headpost outperformed the mask, the difference in precision was 0.03° of visual angle, and the difference in accuracy was less than 0.2°, well within the acceptable range for most use-cases. Our process allows for the production of 3D-printable masks based on CT or MRI scans with the flexibility to incorporate design modifications for experimental equipment and resizing, potentially serving as a promising and more easily accessible alternative to headposts and other NHISs for primate neuroscience research.

## Introduction

Nonhuman primates, like us, use vision as their primary sensory modality to navigate the world, create memories, and make decisions. Given the functional and anatomical similarity between human and nonhuman primate visual systems (Van Essen et al, 2001; Orban et al, 2004; Lafer-Sousa, et al, 2016; Lu et al, 2023), nonhuman primates have long-served as a valuable model for the human visual system. Studies in nonhuman primates enable the execution of experimental paradigms that are not feasible in humans, leveraging the most precise and advanced tools available to neuroscience to measure and perturb neural activity and behavior. While it is possible to carry out some neurophysiology studies in unrestrained monkeys, as is done in certain behavioral studies of monkeys with experimentally induced brain lesions (Vaidya et al., 2018) or those utilizing emerging technologies of wireless neural recording, still very much in their infancy (Dal Monte, et al., 2022; Testard, et al., 2024), the majority of experimental paradigms involving awake, behaving monkeys require that the head be absolutely still. Research that involves eye-tracking and fixation, electrophysiological recording (Wang et al. 2023; Roussy et al. 2022; Hallum et al. 2017; Lemon, 1984), optogenetic perturbation (Shabazi et al., 2024; Azadi et al. 2023b), or functional imaging (Goense, et al., 2010; Lafer-Sousa & Conway, 2013; Tanigawa, et al., 2010) typically necessitate continuous immobilization with minimal head movement for reliable results.

The foundational work of Evarts (1968) and Mountcastle et al. (1975) marked the advent of long-term head fixation techniques. They employed stainless steel bolts anchored in the skull to affix an acrylic head cap. Innovations from various laboratories (Betelak et al. 2001; Evarts 1968;Foeller and Tychsen 2002) have sought to eliminate the acrylic cap. Building on these advancements, Adams et al. (2007) introduced a titanium headpost that is more securely anchored to the skull. This contemporary approach has been rapidly adopted and is now prevalent in most research laboratories. Further surgical innovations have been pursued, such as a low profile halo head fixation system (Isoda et al., 2005; Pigarev, Saalmann, and Vidyasagar 2009; Azimi et al, 2016), but have not been widely adopted.

Despite advancements in headpost design, the approach of surgically attaching a device to the skull inherently carries risks and disadvantages. First, undergoing surgery presents health risks due to anesthesia and postoperative recovery, particularly with respect to infections. Unfortunately, intracranial infection from the headpost is a persistent risk, requiring continual monitoring, cleaning, and maintenance by researchers. Even with this monitoring, headposts can still fail due to infection, necrosis, or bone softening around the headpost and the high torque applied during experiments. Second, these risks are disadvantages to the research project and timeline, because surgery requires extensive physical and personnel resources, cost, and recovery time as the headpost integrates into the bone.

Given these disadvantages and risks, multiple laboratories have independently developed noninvasive approaches to head immobilization. Alternatives have been used in fMRI imaging (Srihasam et al., 2010; Hadj-Bouziane et al., 2014; Drucker et al., 2014), electrophysiology neuronal recordings (Drucker et al., 2014; Amemori et al., 2015) eye fixation tasks (Drucker et al., 2014 Amemori et al., 2015; Machado and Nelson, 2011; De Luna et al., 2014; Rima et al., 2024) and electroencephalographical (EEG) recordings (Nakamura et al., 2020). Most of these head fixation techniques use thermoplastic frames to mold around the precise shape and contours of the monkey’s head to create a customized helmet (Machado and Nelson, 2011; Drucker et al., 2014; De Luna et al., 2014; Slater et al., 2016; Nakamura et al., 2020), which is a similar approach used in human neuroimaging. Most of the research groups lightly sedate the animal for the molding process (Machado and Nelson, 2011; Drucker et al., 2014; Nakamura, et al., 2020), although others create a 3D-printed model of the monkey’s head from MRI scans and use that to mold a thermoplastic frame (Slater et al., 2016), or initially create a mold on a doll’s head and then conduct minor modifications on the awake monkey (De Luna et al., 2014). Other alternative head fixation approaches use negative air pressure (Srihasam et al., 2010; Hadj-Bouziane et al., 2014), aquaplast moldable plastic (Amemori et al., 2015), and 3D-printed chin rests (Rima, et al., 2024).

While there has been a consistent push for noninvasive head fixation techniques, there has not been widespread adoption of the previously described approaches, and headposts remain the predominant method used across research groups. Perhaps this is because the previously mentioned methods still pose some challenges. First, most options use materials that are not commonly used in a primate neuroscience laboratory, such as thermoplastics and plaster, thus requiring both time and training to acquire and learn how to use them. For researchers already familiar with or even trained in surgical approaches, this learning curve can be perceived as cumbersome and time-consuming. Secondly, every mask has to be created anew as monkeys age and grow. These systems thus are wanting for easy reproducibility to accommodate changes in the monkey’s head size over time, presenting an obstacle to consistent long-term use. Third, thermoplastics are shaped by heating and molding, which limits the precision of fine anatomical details, especially for intricate surfaces like those of the face or head. Finally, thermoplastics readily degrade and lose their shape over time.

Our head fixation system uses an emerging technology that is increasingly becoming prevalent in primate neuroscience laboratories or within their larger institutions: the 3D printer. For example, Rima and colleagues (2024) recently used 3D printing to create a chin-rest system. Here, we expand this concept to create a 3D-printed mask that can fully immobilize the monkey’s head. Utilizing CT scans of the monkey’s head and our custom open-source software, FLoRIN (Shahbazi et al. 2018), we engineered a 3D-printable mask system. The approach can also be carried out using anatomical MRI scans. This mask system allows for hi-precision fit, long-term durability, and custom modifications to accommodate experiment-specific needs (e.g. eye-tracking equipment, reward delivery mechanisms, and physiology experiment hardware). The mask can also be readily modified and reprinted to accommodate changes in head shape over time. Similar to Slater et al. (2016), our system comprises a front piece (the mask), which can be used alone for voluntary engagement, and a back piece, which can be attached for full head-immobilization.

After developing this system and demonstrating the feasibility of the production pipeline, we sought to evaluate the efficacy of our mask system for primate neuroscience research. As with the previously discussed alternatives to headposts, we measured our timeline with respect to training monkeys for mask use and then tested whether monkeys wearing masks could match the precision of headposted monkeys on eye fixation tasks. Finally, we report on the success of long-term data collection using our system.

## Methods

All procedures were conducted in accordance with and approved by the National Institute of Mental Health Animal Care and Use and Committee guidelines.

### Subjects

We designed masks for two adult male rhesus macaques named Nd (5 years old, approx. 8 kg) and Gr (8 years old, approx. 14 kg). Nd had previously been implanted with an Opto-Array (Blackrock Neurotech; Rajalingham et al, 2021) over cortical area V4, which had been virally transduced with the depolarizing opsin C1V1, resulting in a cereport pedestal (Blackrock Neurotech) mounted on top of the head, offset from the midline, following procedures similar to those detailed in Azadi et al (2023a). Gr was previously implanted with two chronic multi-electrode Utah Array in the central IT cortex, following similar procedures as in Kar, et al. (2019). This resulted in a custom titanium pedestal housing 4 headstage ports mounted on top of the head, slightly offset from the midline.

Nd and Gr are the two monkeys in our population who had traditional headposts that failed, in part inspiring the development of our mask project. For both monkeys, while they had the headpost, they were trained in head fixation by securing the headpost to their testing chair. Furthermore, they went through our established training protocol to learn to fixate on targets on a computer screen for liquid reward. Nd did not advance far enough in his training to collect eye fixation data. However, while Gr had a headpost, we collected many sessions of data over the course of more than 6 months. Because Gr previously had a headpost and fixation data had been collected, we had the opportunity to conduct a within-subject analysis on precision of eye fixation between a headposted monkey and that same monkey whilst it used the mask.

To test the precision of eye fixation while wearing the mask, we compared Nd and Gr’s eye fixation precision with two other adult males who were similarly trained on an eye fixation task but had a traditional surgically applied headpost, Ph (7 years old, approx. 8.5kg) and Kr (5 years old, approx. 6.8kg). Ph and Kr were implanted with both an Opto-Array over central IT cortex, virally transduced with the depolarizing opsin C1V1, as well as a titanium headpost, again following procedures detailed in Azadi et al. (2023a). This resulted in an acrylic headcap on the midline housing both a cereport pedestal and the caudally located headpost in Kr, and two separate acrylic islands housing the cereport and headpost in Ph.

### Mask Creation

#### Making a 3D base model

The first step of mask creation is the generation of a 3D base model of the monkey’s head. To generate the 3D base model for our mask, we use a structural scan of the monkey’s head, which can be a standard CT or T1-weighted MRI. For Nd and Gr, CT scans were used because the animals had existing implants that were not MR-compatible. For each scan, prior to sedation, animals were premedicated with glycopyrrolate (0.015mg/kg) and ketoprofen (1mg/kg). The animals were then sedated with ketamine (10mg/kg) and dexmedetomidine (0.02mg/kg) and positioned in the scanner. We scanned monkeys on an Epica Vimago HU CT scanner using a 0.2mm resolution available at the NIH Neurophysiology Imaging Facility. Following the scan, monkeys were recovered with revertidine (0.02mg/kg).

We then employed FLoRIN, open course software originally developed for reconstructing neurons and membranes from high-resolution microscopy images (Joesch et al. 2016; Shahbazi 2018, shahbazi et al. 2018), to reconstruct the monkey’s head, creating a precise 3D base model for designing the monkey’s mask, as shown in Fig 1a.

**Figure 1.**
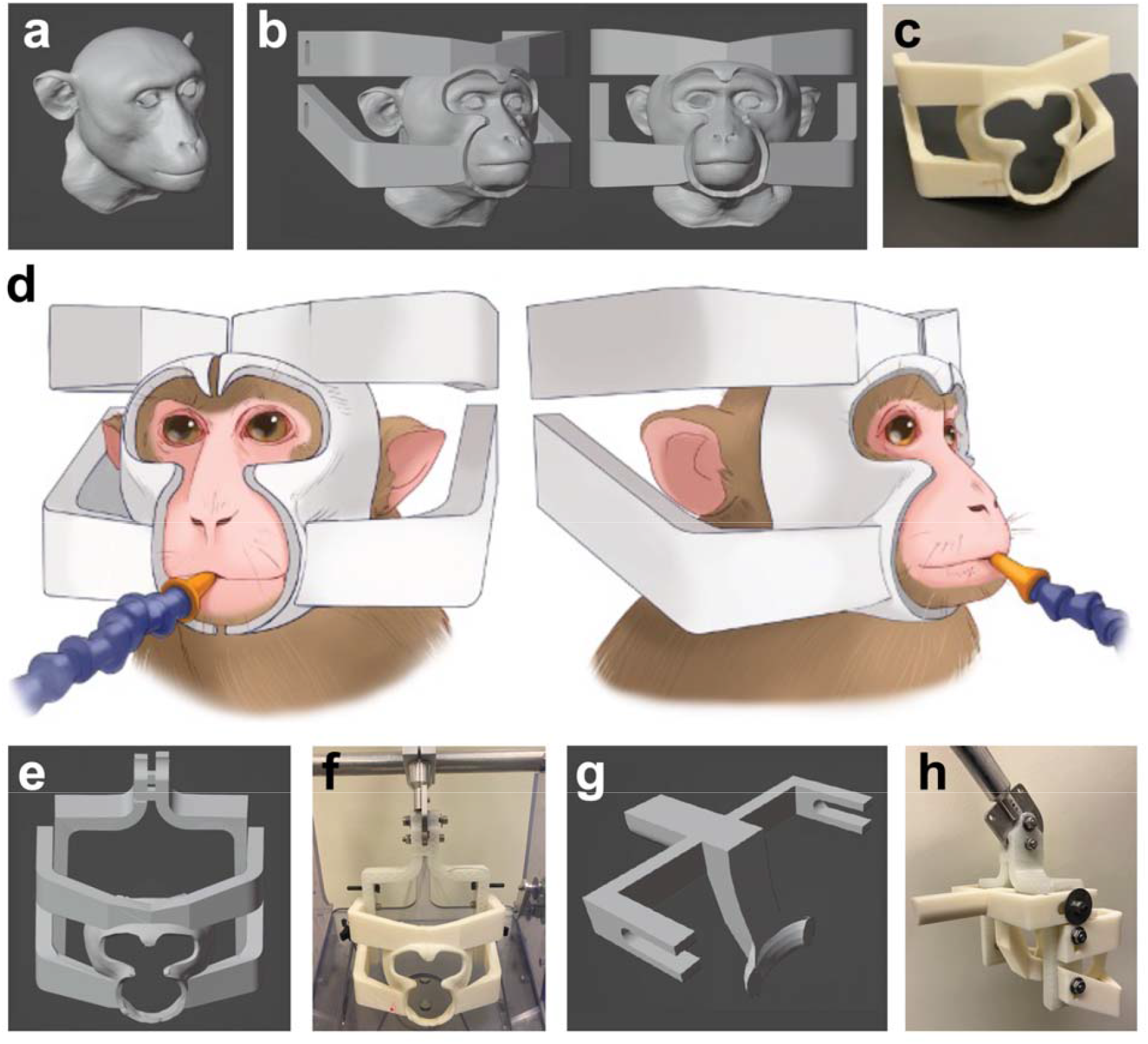
Overview of NHIS Design and Implementation: a) 3D base model (head reconstruction) using FLoRIN based on a standard CT scan of the monkey’s head, providing a detailed and accurate model of the subject’s facial anatomy. b) Design of the NHIS using Blender, illustrating the mask’s structure and various connecting components. c) 3D-printed mask d) Illustrative depiction of the mask in use with a reward delivery system and without the back piece. e) Design of the medial-diapason-shaped connector piece with mask. f) 3D printed mask and medial-diapason connector assembled and mounted on a vertical primate chair (Crist Instrument Co.). g) Design of the back piece for full head-immobilization. h) 3D printed full mask system assembly: includes the mask, back piece, medial-diapason-shaped connector piece, attached to a primate head post holder (Crist Instrument Co.), assembled with nuts, bolts, and washers.

Our custom FLoRIN pipeline is a key innovation in our approach. First, FLoRIN can quickly and easily transform any standard volumetric structural scan into a highly detailed 3D reconstruction of the animal’s head, including the soft tissue structures of the face. Second, FLoRIN’s 3D modeling accurately reflects the current physical state of the monkey’s head, including potential surgical implants. Third, FLoRIN provides compatibility with Python. This, in conjunction with our choice of software for 3D modeling, Blender, allows for the use of Python scripting to convert a series of scan data into 3D models of masks with ease and efficiency.

#### Mask Design

Blender, a software that allows sequences of 3D modeling operations to be exported as code, facilitates mask creation. In Blender, we design a surface with strategically placed cutouts for the eyes, nose, mouth, ears, and top of the head, as well as supports for facial bone structures like the cheekbones, lower jaw, and brow ridge. The designed surface is then conformed to the 3D base model of the monkey’s head using the shrinkwrap modifier, maintaining a default distance of 1mm from the monkey’s face. Following fitting to the base model of the monkey’s head, the mask’s cutouts and supports are then refined and adjusted as needed. The model code can be modified to accommodate diverse experimental hardware devices as needed. In particular, our design leaves the top of the head unobstructed, facilitating the use of commonly implanted devices for optogenetics, electrophysiology, or other cranial chambers and headstages. Figure 1b shows the 3D model of the mask with the 3D base model of the monkey’s head, and Fig 1c shows the 3D printed mask. Figure 1d illustrates the mask in use with a liquid reward delivery system.

In order to attach the mask to the primate testing chair (Crist Instrument Co., Inc.), we used Blender to add connecting components to the 3D model of the mask and to design a 3D-printable medial-diapason-shaped connector (Figure 1e), which mounts on a standard adjustable head post holder (Crist Instrument Co., Inc.). Figure 1f shows the 3D printed mask and medial-diapason-shaped connector attached to the primate chair, which is assembled using nuts, bolts, and washers.

To enable full head immobilization, we used Blender to design a back piece that can attach to the mask from behind and secure the head in the mask (Figure 1g). The back piece design comprises a buttressed curved head support that makes contact along the skull’s occipital crest, as well as a handle and connectors that enable attachment to the mask with nuts, bolts, and washers. During use, a foam pad is attached to the head support and covered with an impermeable disposable shoe cover.

The back piece and connectors are fully customizable based on the monkey’s head position, implants, and chair system. Figure 1h shows the full 3D-printed mask assembly mounted on a standard head post holder (Crist Instrument Co).

#### 3D Printing and Post-processing

For this project, we used the Stratasys F370 3D printer, which offers a print precision of up to 0.127mm (0.005 in.). The material selected for printing was Acrylonitrile Styrene Acrylate (ASA), known for its high durability and versatility in 3D printing applications. ASA exhibits high impact resistance, robust mechanical performance, and strong resistance to UV rays, water, and various chemicals (Carlota 2023). Post-printing, we sanded the mask surface to ensure a smooth surface and double-coated the mask with a resin seal to ensure it could not absorb contaminants and could be properly disinfected between uses. The resin seal also increases durability.

#### Training phase

Once we had a mask ready to introduce to a monkey, we created a training plan to help the monkeys learn to place their head in the mask while seated in their testing chair. The animals were on water control so that we could control the timing and amount of fluid intake. Each monkey’s weight and hydration level were monitored closely and maintained throughout training and data collection. For training, we first desensitized the monkey to the mask, familiarizing them with the object itself by holding the mask in front of the monkey while they were seated in the testing chair. In this session we also paired mask presentation with food and liquid reward to create a positive association. In the following training sessions, we brought the mask closer to the monkey’s face and threaded a straw through the mask or fruit held with long tweezers, so that the monkey would learn to associate moving its head toward the mask with receiving a reward. We then progressed to providing continuous liquid reward while the monkey tolerated affixed mask placement and long-term wear. Within four days, both monkeys willingly engaged with the mask for extended periods to receive juice and water.

The next phase of training was to introduce the back piece, which would allow for full head immobilization (“full mask system”). For this phase, we first placed the affixed mask on the monkey for continuous liquid reward and then progressively desensitized the monkey to the presence of the back piece through to full attachment of the back piece.

After this step, Nd voluntarily (using the mask without the back piece) began maintaining eye fixation on a computer display in exchange for continuous liquid rewards, while Gr immediately began data collection using the back piece. In sum, Nd and Gr required ten and six days of initial exposure to the mask, respectively, before they were sufficiently acclimatized, allowing us to collect data. Figure 2 delineates this behavioral training timeline for mask system stabilization, providing a detailed overview of each acclimatization and training process step.

**Figure 2.**
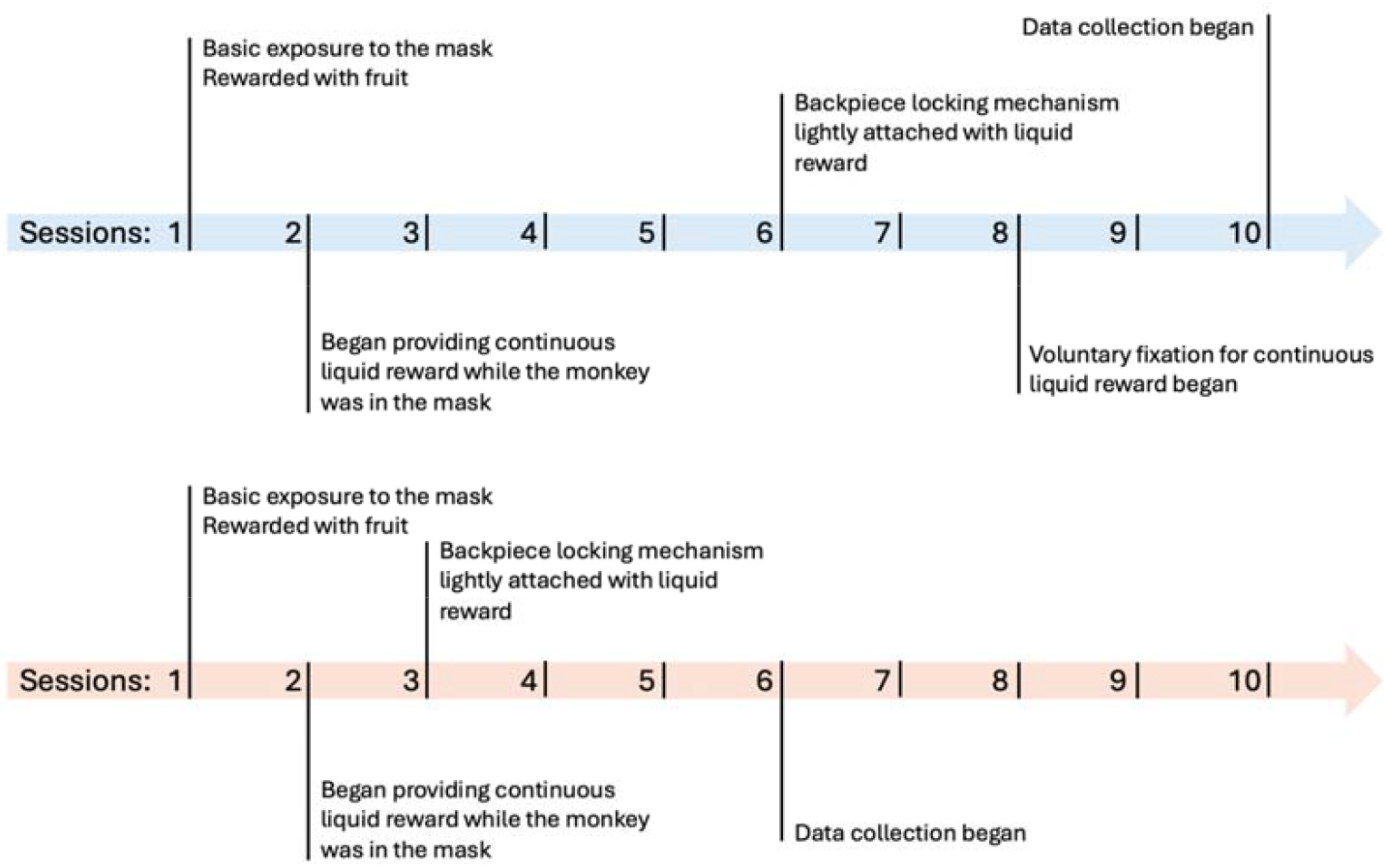
Behavioral training with mask: The training process using our 3D-printed mask system began with familiarizing the monkeys with the mask and employing positive reinforcement for placing their head in the mask. Monkeys were then introduced to the back piece for full head immobilization (in the 6th session for Nd and the 3rd session for Gr). In the 8th session, Nd commenced visual fixation training for continuous rewards without the back piece (voluntary configuration). After two sessions data collection commenced (10th session) in the voluntary configuration (data presented in results section 1). Subsequently, data collection with the full mask system began (data presented in results section 2). Gr immediately began data collection with the full mask system (6th session, data presented in results section 2). The timelines for Nd and Gr are depicted in blue and orange, respectively.

### Data Collection and Analysis: Comparison with Traditional Headpost

#### Voluntary Head Immobilization with Mask System vs Traditional Headpost

We first conducted a comparative analysis of visual fixation data (eye-tracking) using two distinct restraint systems on two monkeys: one with a traditional headpost (Monkey Ph) and the other using our non-invasive mask-based system, without the back piece (Monkey Nd). Crucially, entry into the mask immobilization system was entirely voluntary for monkey Nd—at no point during this session was the locking back piece of the mask engaged. This aspect of the study design was implemented to assess the effectiveness and compliance of the mask-based system under its least restrictive condition. These data were collected in Nd as soon as initial training with the mask system was complete (see Figure 2 training plot).

The monkeys performed a simple 13-target fixation task. The animals were positioned 57 cm from a calibrated 32” screen (120 Hz, 1920×1080 IPS LCD, Cambridge Research System Ltd). In each trial, a fixation target (0.2 degrees in size) was presented for 500 ms seconds at one of 13 locations on the computer display (Figure 3a). Target locations were uniformly distributed across the screen (Figure 3a). The most peripheral target location was approximately 14 degrees from the visual field center (10 degrees away both horizontally and vertically). The monkeys were tasked with continuously fixating on each target during the presentation window. Compliance was reinforced through liquid rewards via a sipper tube. Data were collected using custom NIMH MonkeyLogic scripts on a Dell computer. Eye tracking was carried out with an Eyelink 1000 Plus (SR Research), at a frequency 1000 Hz.

**Figure 3.**
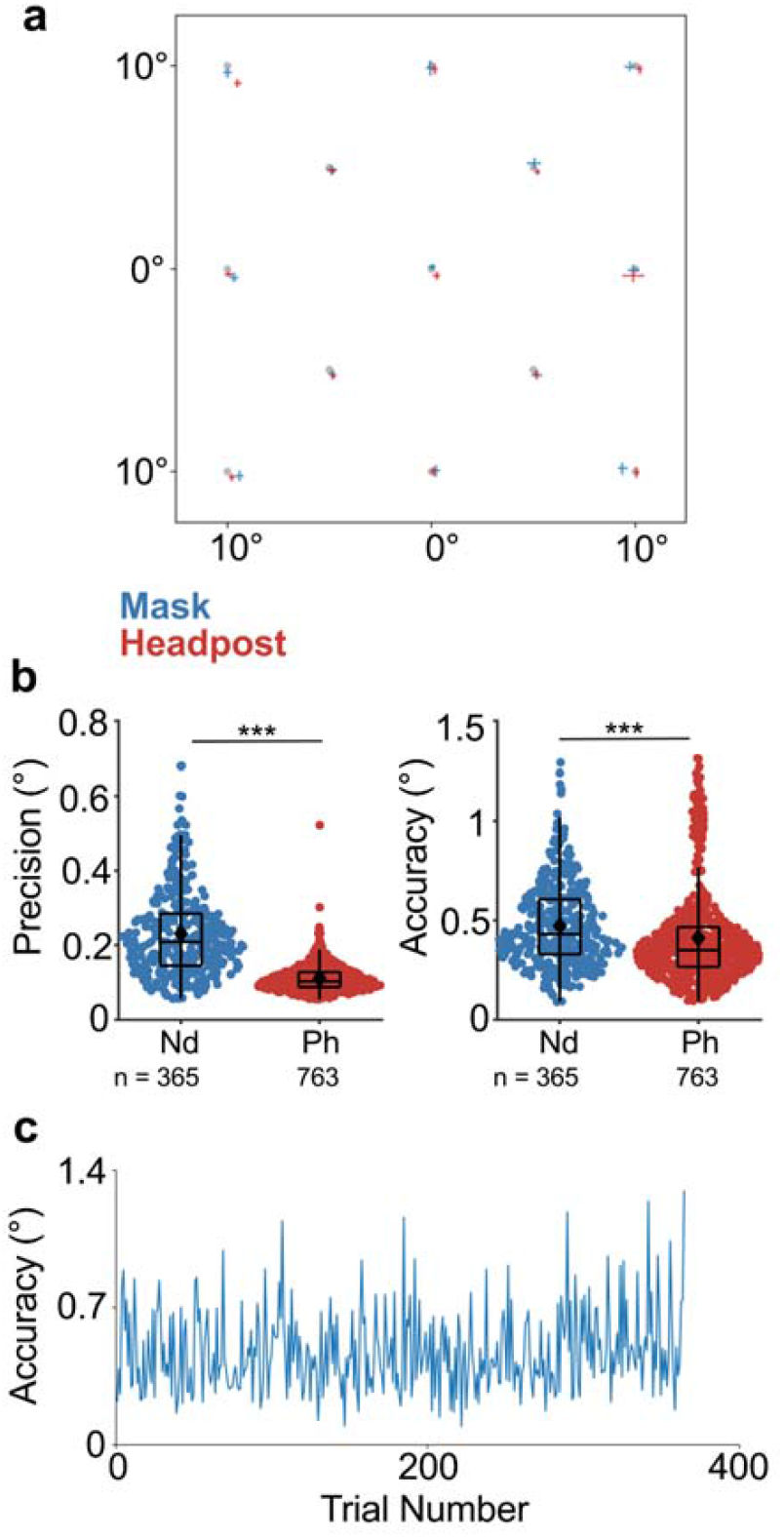
Accuracy and precision of visual fixation eye-tracking data collected using voluntary head-immobilization configuration of the mask NHIS vs traditional headpost: 2 monkeys (Nd, mask-restrained, blue; and Ph, headposted, red) fixated on 13 targets throughout the testing procedure, receiving a liquid reward, recorded using an eye tracker. a) Fixation targets and mean fixation positions for monkeys Nd and Ph. X-axis and y-axis show horizontal and vertical degrees of visual angle on the computer screen, and error bars show ±1 SD. b) Trial-by-trial gaze precision and accuracy for monkeys Nd (n = 365 trials) and Ph (n = 763) during the 13-target fixation task. Y-axes show mean gaze precision and accuracy in degrees of visual angle for the left and right plots, respectively. Means are indicated with a black marker. Black outlines show boxplot statistics: boxes represent interquartile range (IQR) with median, and whiskers represent 1.5*IQR from the quartiles. The Mann-Whitney U test indicated a significant difference in gaze precision between the mask (median = 0.2010°, IQR = 0.1402°) and headpost (median = 0.1029°, IQR = 0.0403°) conditions, U = 32,229, p < 0.001. Similarly, the test revealed a significant difference in gaze accuracy between the mask (median = 0.43°, IQR = 0.2765°) and headpost (median = 0.3501°, IQR = 0.2000°) conditions, U = 93,744, p < 0.001. Precision and accuracy were significantly better in the headpost condition compared to the mask condition. c) Gaze accuracy for monkey Nd across trials in 13-target fixation task. Y-axis shows gaze accuracy in degrees of visual angle, and x-axis shows trial number. Linear regression showed no significant correlation between trial index and gaze accuracy for Nd (slope = 0.00019, R^2^ = 0.0100, p = 0.55).

To assess the efficacy of our voluntary mask immobilization system compared to the traditional headpost method, we compared eye tracking precision and accuracy of gaze fixations (Figure 3b). Both values were calculated on a per-trial basis: precision was defined as the mean distance (in degrees of visual angle) between each gaze position sample and the mean gaze position for that trial, and accuracy was defined as the mean distance (in degrees of visual angle) between each gaze position sample and the fixation target for that trial. Eye tracking samples were taken from the 500ms target fixation period in each trial, and trials in which monkeys broke fixation were not taken into account. We then performed a Mann-Whitney U test to compare the distributions of Nd and Ph’s trial-by-trial gaze precisions, as well as their trial-by-trial gaze accuracies. This non-parametric test was chosen because the data did not meet the assumptions of normality (Shapiro-Wilk test), and the two datasets had unequal sample sizes. The test evaluates whether the distributions of gaze metrics for the two conditions are significantly different.

Next we assessed the stability of gaze accuracy over the course of the session while Nd was using the mask in the voluntary configuration. This analysis was important because, in the voluntary configuration, Nd could pull his face back at any time during the session to take a break, and then place his face back in the mask to continue data collection without recalibration. We wanted to know whether such disengagement and reengagement events impacted our gaze accuracy measurements. We fit a linear model to Nd’s gaze accuracy data to determine whether there was any correlation between trial index and Nd’s trial-by-trial gaze accuracy. A positive correlation would indicate increasing instability over the course of a session, while negative or no correlation would indicate stable fixation across the session.

#### Full Immobilization with Mask System vs Traditional Headpost

In order to test the relative efficacy of the full-immobilization mode of the mask system (back piece engaged) versus a standard headpost in a typical experimental setup, we analyzed fixation data acquired in two monkeys using the full-immobilization mask system (Nd & Gr) and three monkeys with traditional headposts (Ph, Gr, & Kr) while they performed a cortical perturbation detection (CPD) task (Azadi et al., 2023b), as part of ongoing research projects.

Note that Gr was tested with both immobilization systems (mask and headpost), enabling a within-subject comparison. Gr, being a particularly large animal (14 kg), also provided a robust test of the mask system’s durability. At this stage of data collection, all four monkeys had been trained to perform the CPD task, with two of the monkeys (Ph & Kr) tasked with detecting optogenetic stimulation delivered to central IT, one monkey (Nd) tasked with detecting optogenetic stimulation delivered to visual area V4, and the fourth monkey (Gr) tasked with detecting electrical microstimulation delivered to central IT. Each trial in the CPD task was initiated when the monkey fixated a central fixation target (0.4 degrees in diameter) for 500 ms. Then, an image was displayed for 1 second, during which, in half of the trials randomly selected, cortical stimulation was delivered for 200 ms halfway through the image presentation. After the image disappeared, the monkey was rewarded for correctly identifying whether the trial contained cortical stimulation by saccading to one of two targets at the top and bottom of the screen. For the purposes of the present study, fixation data were extracted from a random selection of these CPD task sessions, which were collected over many months. For each monkey, five sessions were analyzed.

We compared the efficacy of our mask system in its full-immobilization configuration against the headpost during long-term data collection while monkeys performed the CPD task, by analyzing the trial-by-trial gaze precision and accuracy of fixations (Figure 4). We considered gaze position samples collected during the image presentation phase of each trial, throughout which central fixation was required. Trials in which fixation was broken at any point after trial initiation were not taken into account. Precision and accuracy were calculated using the same method as above.,. For each gaze fixation metric (precision and accuracy), we performed a linear mixed-effects model analysis using MATLAB’s fitlme function. The model included a fixed effect for condition (mask or headpost) and a random intercept for subject (monkey) to account for between-subject variability. The headpost condition was set as the reference category. The gaze fixation metric (either precision or accuracy) for each trial was used as the dependent variable, and we tested for significance using t-statistics and p-values for the fixed effects. Model coefficients were estimated using restricted maximum likelihood (REML). The same analysis was performed for gaze precision and gaze accuracy. This approach allowed us to examine the difference in gaze fixation metrics between the mask and headpost conditions, while controlling for individual differences in baseline gaze fixation.

**Figure 4.**
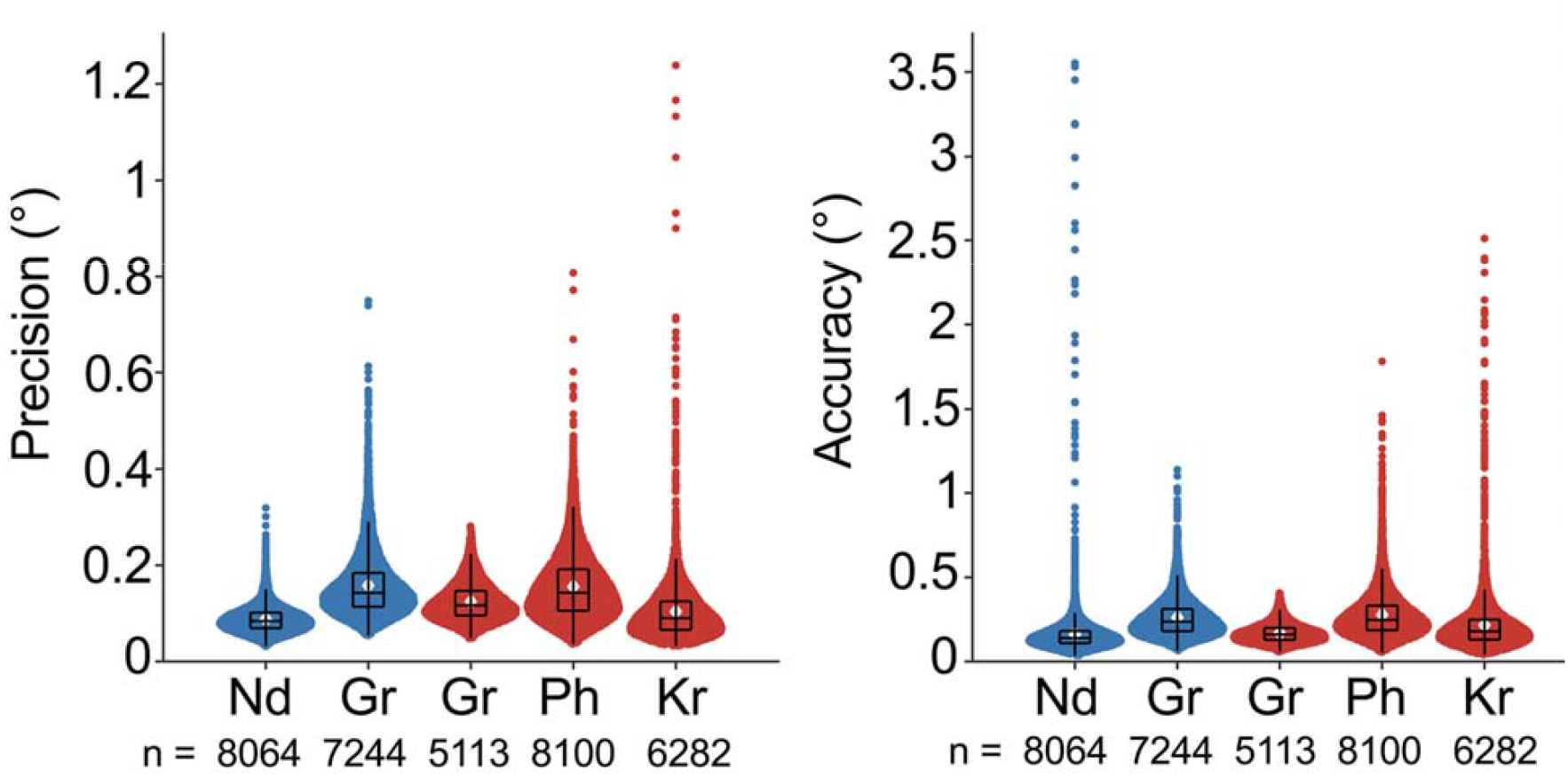
Accuracy and precision of visual fixation eye-tracking data using full head-immobilization configuration of the mask NHIS vs traditional headpost. Individual monkeys’ trial-by-trial gaze precision (left) and accuracy (right) distributions during central fixation in a cortical perturbation detection (CPD) task. Data collected in monkeys head-immobilized with the mask NHIS (Nd, Gr) are shown in blue, and those head-immobilized with a headpost (Gr, Ph, Kr) are shown in red. Y-axis shows gaze precision (left) and accuracy (right) in degrees of visual angle. Means are indicated with a white marker. Black outlines show boxplot statistics: boxes represent interquartile range (IQR) with median, and whiskers represent 1.5*IQR from the quartiles. Data shown for all plots is from n= 34803 trials (Nd: 8064; Gr with mask: 7244; Gr with headpost: 5113; Ph: 8100; Kr: 6282). A linear mixed-effects model revealed a significant effect of head stabilization method on gaze precision and accuracy. The mean gaze precision in the headpost condition (the reference condition) was estimated to be 0.1096° (SE = 0.0184°), t(34,801) = 5.94, p < 0.001. Gaze precision in the mask condition was significantly higher (worse) compared to the Headpost condition, with a difference of 0.0337° (SE = 0.0010°), t(34,801) = 32.52, p < 0.001. The mean gaze accuracy in the headpost condition was estimated at 0.1832° (SE = 0.0379°), t(34,801) = 4.83, p < 0.001. Gaze accuracy in the mask condition was significantly higher than in the headpost condition, with an estimated difference of 0.0914° (SE = 0.0024°), t(34,801) = 37.73, p < 0.001. A Mann-Whitney U test was used to carry out within-subject pairwise comparisons between conditions for Gr, for both gaze precision and accuracy. The results indicated a significant difference in gaze precision between the mask (median = 0.143°, IQR = 0.070°) and headpost (median = 0.117°, IQR = 0.051°) conditions, U = 12,331,258, p < 0.001. The test revealed a significant difference in gaze accuracy between the mask (median = 0.237°, IQR = 0.133°) and headpost (median = 0.161°, IQR = 0.072°) conditions, U = 8,252,961, p < 0.001. Precision and accuracy were significantly better in the headpost condition compared to the mask condition.

Because we collected data in monkey Gr using both the mask and headpost head-immobilization methods, we were able to carry out a within-subject analysis. We used a Mann-Whitney U test to compare the distribution of trial-by-trial gaze precisions, as well as the trial-by-trial gaze accuracies, between the mask and headpost conditions. This non-parametric test was chosen because the data did not meet the assumptions of normality (Shapiro-Wilk test), and the two conditions had unequal sample sizes. The test evaluates whether the distributions of gaze metrics for the two conditions are significantly different.

## Results

### Voluntary Head Immobilization with Mask System vs Traditional Headpost

In the initial comparative analysis between a traditional headpost and our mask system without the back piece engaged, Nd (voluntarily mask-immobilized) completed 365 trials over 1 session and Ph (headposted) completed 673 trials over 1 session. Figure 3 shows the trial-by-trial distributions of the two monkeys’ gaze precision and accuracy. The difference in mean gaze precision between Nd and Ph was 0.12° of visual angle, and the difference in mean gaze accuracy was 0.060° (Nd measurement - Ph measurement). These results indicate that the precision and accuracy of gaze fixations were slightly better in the headpost condition. While these differences were small in magnitude, they were significant. The Mann-Whitney U test indicated a significant difference in gaze precision between the mask (median = 0.2010°, IQR = 0.1402°) and headpost (median = 0.1029°, IQR = 0.0403°) conditions, U = 32,229, p < 0.001. Similarly, the test revealed a significant difference in gaze accuracy between the mask (median = 0.43° degrees, IQR = 0.2765°) and headpost (median = 0.3501°, IQR = 0.2000°) conditions, U = 93,744, p < 0.001. Overall, the results show a statistically significant but slight decrease in eye tracking performance for Nd (mask-stabilized) compared to Ph (headpost-stabilized).

In the voluntary configuration, Nd could pull his face back from the mask at any time during the session, and then return his face to the mask to continue data collection. We did not recalibrate the eye-tracker between these events. In order to assess the impact of such disengagement and re-engagement events, we analyzed Nd’s gaze accuracy over time and found it to be consistent throughout the session (Figure 3c): Linear regression showed no significant correlation between trial index and Nd’s trial-by-trial gaze accuracy (slope = 0.00019, R^2^ = 0.0100, p = 0.55). These results show that disengagement and re-engagement events did not impact our gaze accuracy measurements.

### Full Immobilization with Mask System vs Traditional Headpost

In the subsequent analysis comparing standard headposts and the full-immobilization mode of the mask system (front and back piece engaged) during long-term data collection in a typical experimental set-up (Figure 4), across the 5 sessions per monkey that we analyzed, Nd (fully mask-immobilized) completed 8064 trials, Ph (headposted) completed 8100 trials, and Kr (headposted) completed 6282 trials. Gr, the within-subject comparison monkey, completed 7244 trials across 5 sessions while fully mask-immobilized, and completed 5113 trials across another 5 sessions while headposted. On average, monkeys who were immobilized using our mask system completed more trials than monkeys who were headposted. Additionally, long-term data collection using the mask was successful; over 8 months, Nd completed approximately 100,000 trials, and Gr completed 30,000 trials over the course of 3 months while using the mask.

Linear mixed-effects models revealed small but significant effects of head-stabilization method on gaze precision and accuracy. The mean gaze precision in the headpost condition was estimated to be 0.1096° (SE = 0.0184°), t(34,801) = 5.94, p < 0.001. Gaze precision in the mask condition was significantly worse compared to the headpost condition, with a difference of 0.0337° (SE = 0.0010°), t(34,801) = 32.52, p < 0.001. The 95% confidence interval for the effect of the mask condition was [0.0317°, 0.0358°]. The average gaze accuracy in the headpost condition was estimated at 0.1832° (SE = 0.0379°), t(34,801) = 4.83, p < 0.001. Gaze accuracy in the mask condition was significantly worse than in the headpost condition, with an estimated difference of 0.0914° (SE = 0.0024°), t(34,801) = 37.73, p < 0.001. The 95% confidence interval for the effect of the Mask condition was [0.0866°, 0.0961°].

For our within-subject comparison in Gr, there was a difference of 0.033° in mean gaze precision and a 0.089° difference in mean gaze accuracy when using the mask compared to a traditional headpost (mask measurement - headpost measurement). While these differences were small in magnitude, they were significant. The Mann-Whitney U test indicated a significant difference in gaze precision between the mask (median = 0.143°, IQR = 0.070°) and headpost (median = 0.117°, IQR = 0.051°) conditions, U = 12,331,258, p < 0.001. Similarly, the test revealed a significant difference in gaze accuracy between the mask (median = 0.237°, IQR = 0.133°) and headpost (median = 0.161°, IQR = 0.072°) conditions, U = 8,252,961, p < 0.001. Precision and accuracy were significantly better in the headpost condition compared to the mask condition.

## Discussion

In our study, monkeys with their heads immobilized using a 3D-printed mask demonstrated eye-tracking precision and accuracy similar to that of their headposted counterparts. While headposted monkeys demonstrated significantly better gaze precision and accuracy during fixation, the difference observed when using the NHIS in its least restrictive configuration was just 0.12° for precision and 0.06° for accuracy (pairwise comparison), and in its most restrictive configuration was 0.03° for precision and 0.18° for accuracy (mixed-effects model estimate). In our within-subject comparison, the differences were similar: 0.03° for precision and 0.09° for accuracy. Though statistically significant, this minimal loss in gaze precision and accuracy using our NHIS relative to the traditional headpost is well-within the acceptable range for most use cases. Typically, experiments carried out with head-fixed awake behaving monkeys require monkeys to maintain fixation within a not less than 0.8° x 0.8° window, including studies of V1 neurons whose receptive fields tend to be small (Abhishek & Horwitz, 2022). Thus, the quality of eye-tracking with NHIS head fixation is comfortably within the range of suitability for most experimental needs.

Moreover, in this study, we used Eyelink 1000 (SR Research Ltd.), which has a precision of 0.01º and accuracy down to 0.15º for tracking and recording gaze position under ideal conditions. Most other video-based eye trackers commonly used in nonhuman primate physiology experiments do not offer such high precision and accuracy. The performance of the mask introduced in this study is comparable to, or even exceeds, some other popular systems used in similar research. For example, the ViewPoint EyeTracker (Arrington Research, Inc.) has a precision of 0.15º and an accuracy ranging from 0.25º to 1.0º.

Furthermore, monkeys stabilized with the mask were able to perform as many if not more trials than those stabilized using a headpost. As well, the monkeys were able to complete months of data collection using the mask, demonstrating long-term suitability of the mask and the continual cooperation of monkeys to engage with the mask. As to whether our NHIS can remain stable with larger and stronger monkeys, we successfully used our NHIS with a relatively large adult male who was 14kg. There were no complications from his use of the NHIS for 30,000 trials over 3 months, thus demonstrating the mechanical robustness of our system. The comparable performance observed between headposted monkeys and ones wearing our mask comes with a range of benefits. First, reducing the need for surgery and using a non-invasive approach for head immobilization aligns with the continual mission to refine methods and minimize the potential for pain and stress in research animals, as outlined in the principles of the 3Rs (Russell, Burch, and Hume, 1959). While the tools commonly used to measure and perturb neural activity and behavior still require surgical interventions, the field is moving towards the use of wireless devices (Dal Monte, et al., 2022; Testard, et al., 2024) which will reduce the persistent risk of infection posed by implants that breach the skin barrier. Replacing headposts with effective non-invasive head-immobilization systems goes hand-in-hand with these efforts. Moreover, reducing the need for headpost placement surgery and recovery allows for a faster transition to data collection. The mask system can be quickly produced in-house using 3D printing technology, and the monkeys in our study learned to use the mask within 10 days.

Furthermore, the use of 3D printing to produce an NHIS, as presented here, offers several advantages over the commonly used thermoplastic materials in most NHIS designs (Machado and Nelson, 2011; Drucker et al., 2014; De Luna et al., 2014; Slater et al., 2016; Nakamura et al., 2020). First, 3D printing allows for precise anatomical fit, better accommodating unique individual differences than thermoplastics can. Thermoplastics are shaped by heating and molding, limiting the precision of fine anatomical details, especially for intricate surfaces like the face or head. Second, designing the mask in a 3D modeling software environment allows for complex geometries that might improve comfort, stability, and fit beyond the capabilities of molded thermoplastic. Third, such an approach enables design modifications to suit specific experimental needs and equipment configurations (e.g., cranial implants, style of primate chair, reward delivery system), enhancing overall utility, integration, and production. For example, the design presented here affords additional space on the top of the head for the placement of research-related implants, in contrast to full-head thermoplastic helmets. Fourth, 3D-printed materials can be designed to withstand repeated use and cleaning, potentially outlasting thermoplastic molds, which may degrade or lose shape over time. Finally, the mask can be readily modified and re-printed without sedation or excessive downtime to accommodate changes in head size due to animal growth, experimental needs, or to replace damaged parts.

Our assessment of the mask system, as presented in this paper, has some potential limitations. The sample was restricted to two non-naive monkeys, both of which had already undergone fixation training. We anticipate naive monkeys will take longer to acclimate to the mask, possibly extending the training process beyond the timeline we have outlined here (Fig 2). Also, the efficacy of our system in providing stable intracranial recordings is yet to be shown. We hope that sharing this method with the greater scientific community brings interest to shed more light on these unaddressed aspects. In addition, the mask creation pipeline requires sedation, access to an anatomical scanner (CT or MRI), and an overnight or longer 3D printing process, as well as post-processing, before use. Future design iterations may minimize the post-processing required, but the speed of available 3D printing technology and requisite sedated scanning present a ceiling on the extent to which the mask system can be streamlined. Finally, the scope of the present work does not evaluate the mask’s suitability for acute electrophysiology or functional imaging, as has been done for other NHIS systems (Srihasam, et al., 2010; Amemori, et al., 2015; Drucker, et al., 2015; Slater, et al., 2016). We expect that our system will perform at least as well as those systems, but further testing is needed.

Going forward, we plan to continue to monitor the efficacy of the mask system for data collection, as well as incorporate and test additional options and functionality. For example, our in-house CT scanner allows us to access scans for many monkeys, so we are currently developing and testing an option to have a mask design based on average macaque 3D base models (averaged within sex). This standardized ready-made mask would be suited for use in experiments where stringent immobilization is not required. The average face mask could be used and scaled to accommodate growth and slight changes in monkeys’ face structure without requiring sedation or scanning. We are also testing a double-layer design to allow for a softer, padded material to form the interior surface of the mask to increase monkeys’ comfort. Finally, there is potential for our NHIS mask, without the back piece, to be placed in the front of the monkey’s home cage, with cage modifications or other approaches. The opportunity to collect precise eye-tracking data in an unrestrained monkey trained to place its head in the mask, unobstructed by cage bars from the eye-tracker or stimuli, would be an exciting potential to further refine our experimental approaches for neuroscience research in awake and behaving monkeys.

## Availability statement

The production pipeline code will be made publically available to encourage widespread adoption.

## Acknowledgments

We thank Dr. Alex Cummins and Katherine Cameron for their crucial contributions to 3D printing. Anatomical scanning was carried out in the Neurophysiology Imaging Facility Core (NIMH, NINDS, NEI). This research was supported by the Intramural Research Program of the NIMH ZIAMH002958 (to A.A.)

## Notes

### Competing Interest Statement

The authors have declared no competing interest.

### Summary of Updates

The manuscript has been fully updated. Two new subjects have been added, and the data and results have been updated according to the new data.

