## Supplementary material for "3D-Printable Non-invasive Head Immobilization System for Non-Human Primates": Supplimentry Materials

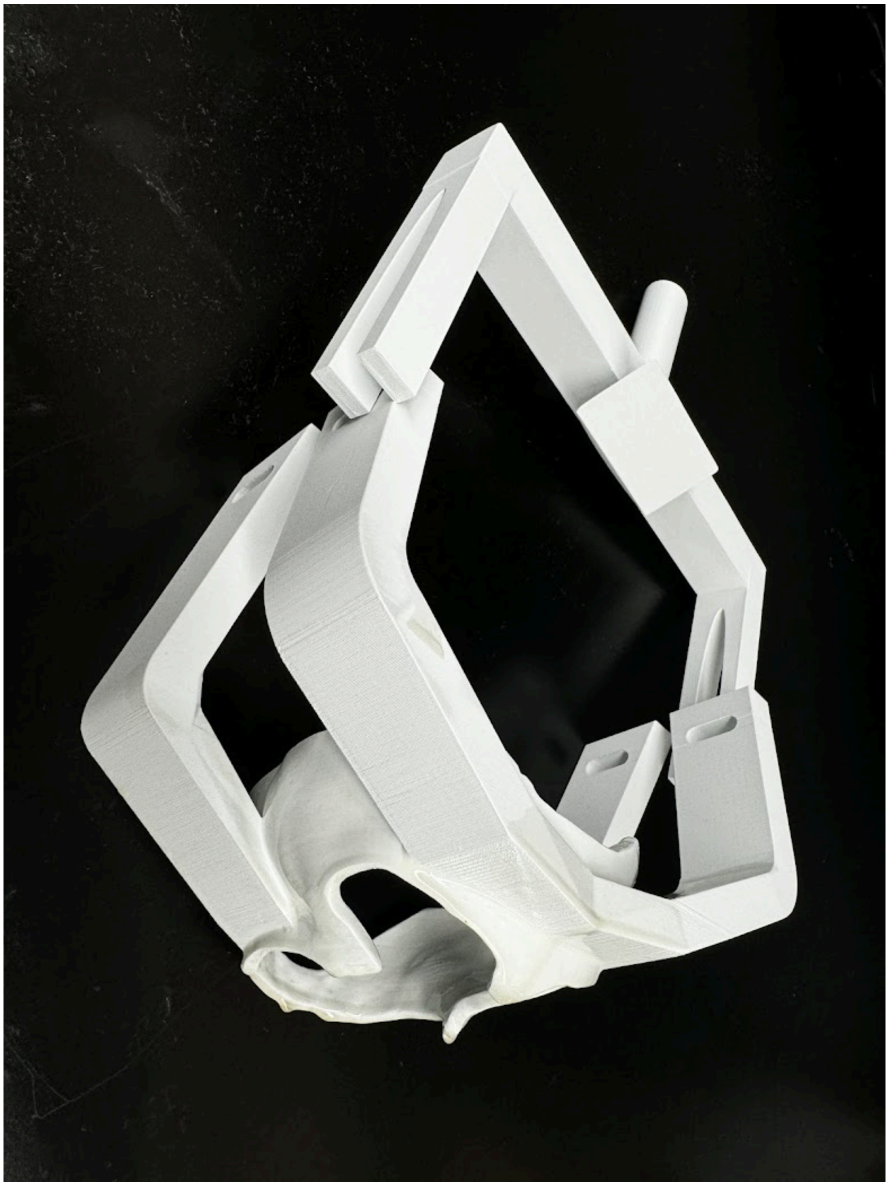

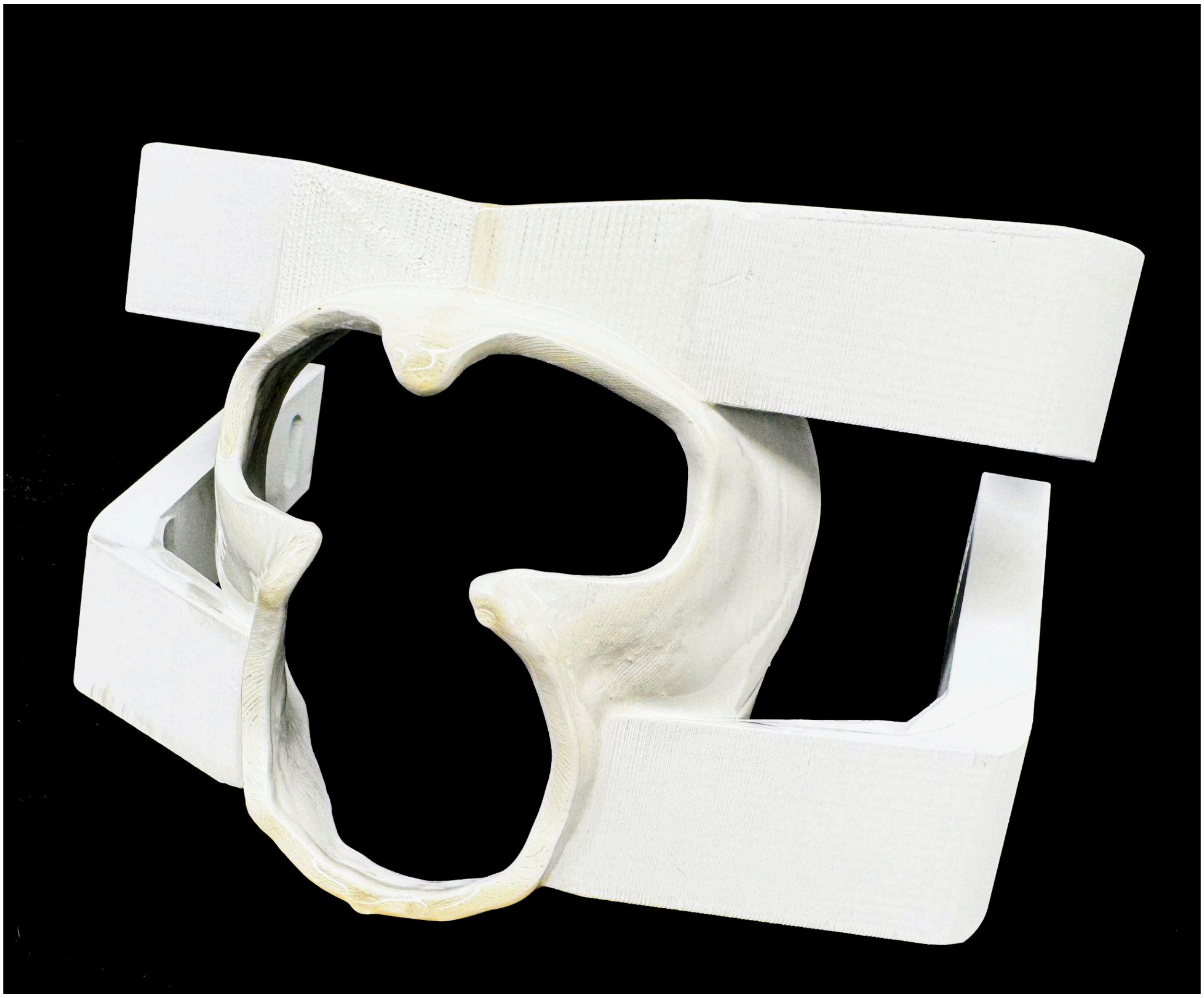

Sup-fig 1: 3D printed mask with backpiece.

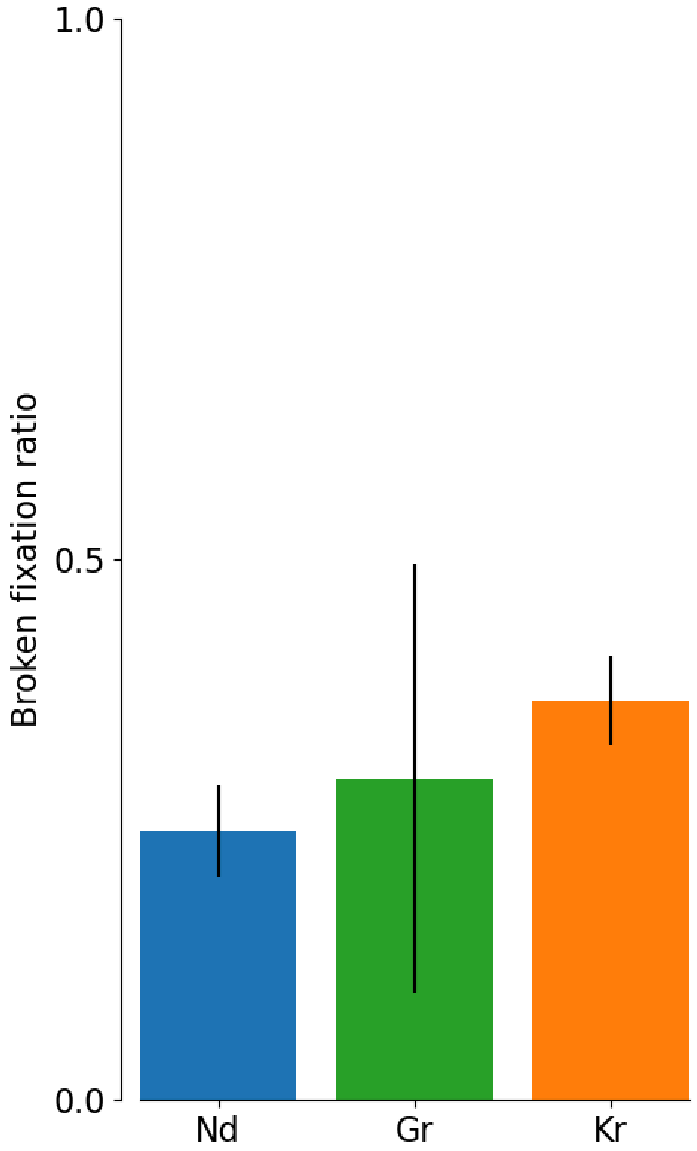


Figure SX. Broken Fixation Ratios During Task. Bars show the proportion of trials during which monkeys Nd (NHIS), Gr (NHIS), and Kr (Headpost) broke fixation. Error bars show 1 SD and data from n=21718 trials across five sessions.(Blue: Nd, Green Gr and Orange is Kr respectively)

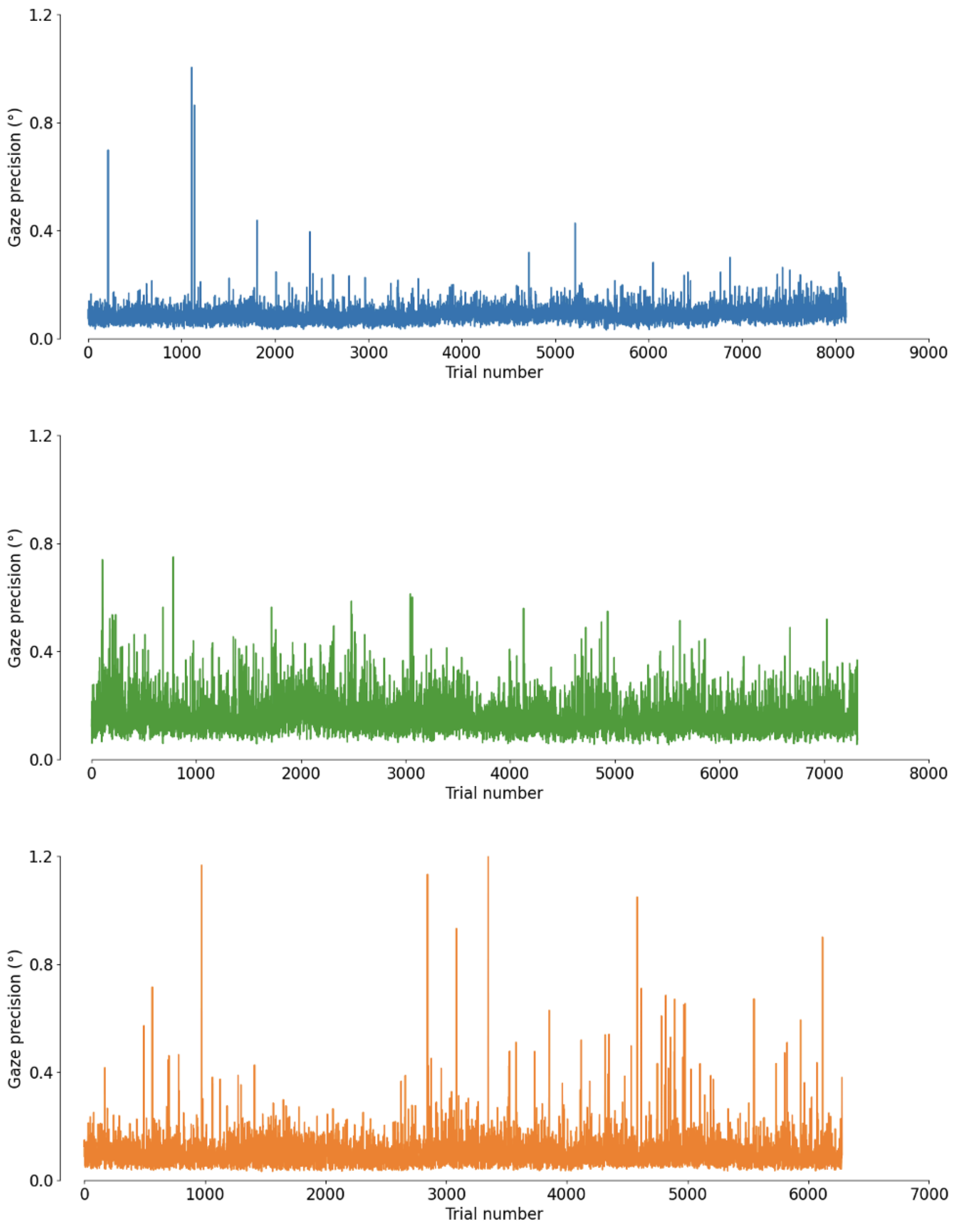


Figure SX. Gaze Precision During Task. Y-axes show the precision of gaze in degrees for monkeys Nd (NHIS, top), Gr (NHIS, middle), and Kr (headpost, bottom); x-axes show trial number. Data shown from n=21718 trials. (Blue: Nd, Green Gr and Orange is Kr respectively)

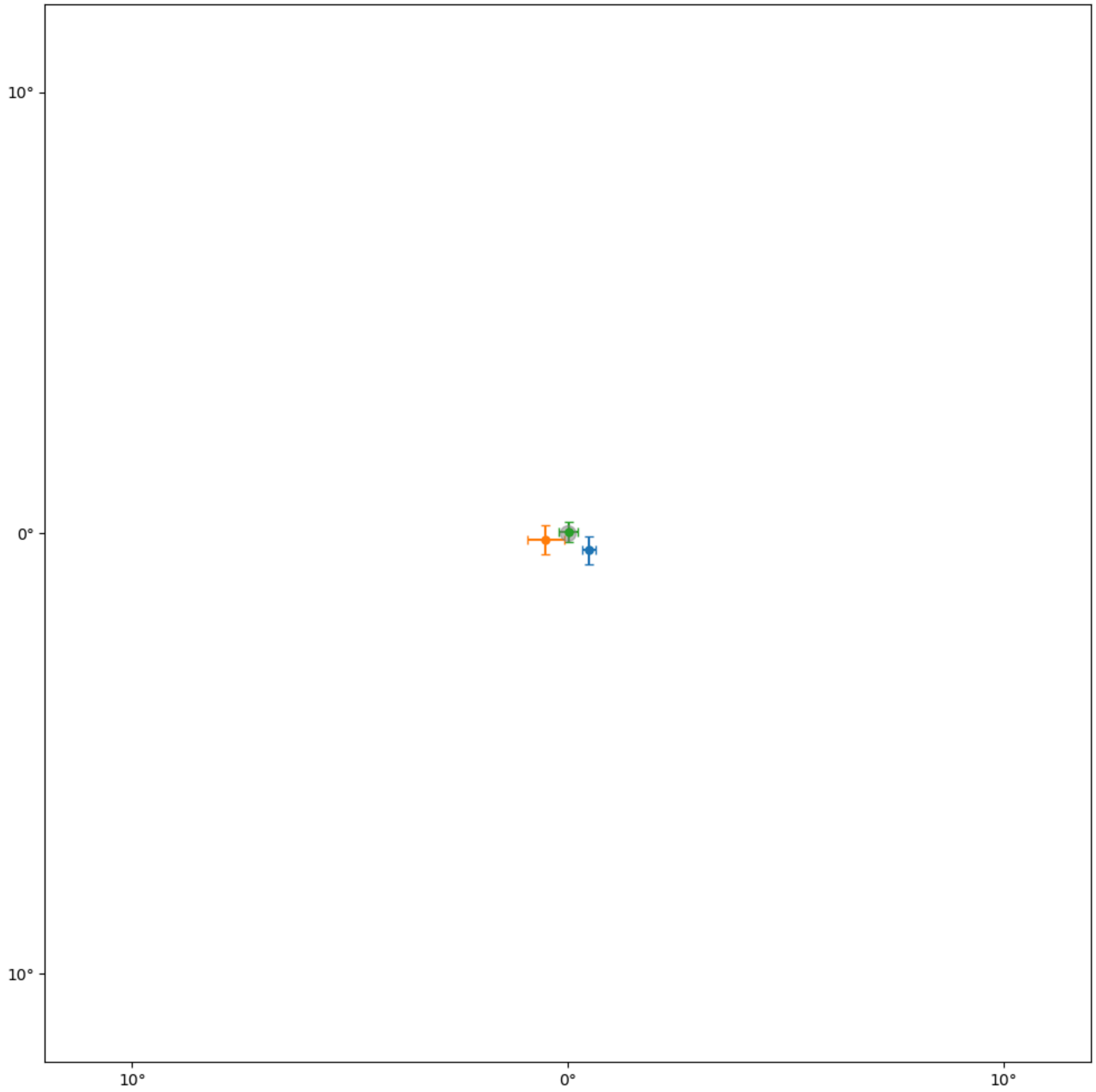


Figure SX. Fixation During Task. Grey dot shows fixation target, points show mean fixation positions for monkeys Nd (NHIS, blue), Gr (NHIS, green), and Kr (NHIS, orange) during central fixation in a 2AFC task. Error bars show 1 SD. Data shown from n=21718 trials.
